# Autonomic/Central Coupling Boosts Working Memory in Healthy Young Adults

**DOI:** 10.1101/2020.04.22.056481

**Authors:** Pin-Chun Chen, Lauren N. Whitehurst, Mohsen Naji, Sara C. Mednick

**Author notes:** **Corresponding author:** Sara C. Mednick, PhD,Department of Cognitive Science University of California, Irvine, Irvine, CA 92697. **Grant support:** NIH R01AG046646; Office of Naval Research, Young Investigator Award to Mednick. **Competing interests:** The author(s) declare no competing interests.

## Abstract

Working memory (WM) is an executive function that can improve with training. However, the precise mechanism for this improvement is not known. Studies have shown greater WM gains after a period of sleep than a similar period of wake (Kuriyama et al. 2008a; Zinke, Noack, and Born 2018), with WM improvement correlated with slow wave activity (SWA; 0.5-1Hz) during slow wave sleep (SWS) (Sattari et al. 2019; Pugin et al. 2015; Ferrarelli et al. 2019). A different body of literature has suggested an important role for autonomic activity during wake for WM (Hansen et al. 2004; Mosley, Laborde, and Kavanagh 2018). A recent study from our group reported that the temporal coupling of autonomic and central events (ACEs) during sleep was associated with memory consolidation (Naji et al. 2019). We found that heart rate bursts (HR bursts) during non-rapid eye movement (NREM) sleep are accompanied by increases in SWA and sigma (12-15Hz) power, as well as increases in the high-frequency (HF) component of the RR interval, reflecting vagal rebound. In addition, ACEs predict long-term, episodic memory improvement. Building on these previous results, we examined whether ACEs may also contribute to gains in WM. We tested 104 young adults in an operation span task (OSPAN) in the morning and evening, with either a nap (with electroencephalography (EEG) and electrocardiography (ECG)) or wake between testing sessions. We identified HR bursts in the ECG and replicated the increases in SWA and sigma prior to peak of the HR burst, as well as vagal rebound after the peak. Furthermore, we showed sleep-dependent WM improvement, which was predicted by ACE activity. Using regression analyses, we discovered that significantly more variance in WM improvement could be explained with ACE variables than with overall sleep activity not time-locked with ECG. These results provide the first evidence that coordinated autonomic and central events play a significant role in sleep-related WM improvement and implicate the potential of autonomic interventions during sleep for cognitive enhancement.

## 1. Introduction

Working memory (WM), the ability to retain, manipulate, and update information over short periods of time, is essential to higher order cognition (e.g. language comprehension, reasoning, and general intelligence; Engle and Kane 2004) and for performing many daily activities (Cantarella et al. 2017; Kane et al. 2007). WM is prone to age-related cognitive decline, which is evident in midlife, and becomes particularly pronounced in older age (age > 75) (Hale et al. 2011). Considering the importance of WM for cognitive functions and the detrimental effects of age-related WM declines, the possibility of improving WM performance to facilitate cognitive health has been advanced (Zinke et al. 2013; Soveri et al. 2017). Importantly, recent studies suggest that WM capacity is subject to experience-dependent change (Au et al. 2015; Jaeggi et al. 2008; Karbach and Verhaeghen 2014), but the mechanism underlying this change is not understood. Recent studies suggest a role for sleep (Sattari et al. 2019; Ferrarelli et al. 2019; Pugin et al. 2015). The purpose of the current study is to identify sleep-specific features that support WM enhancement.

Sleep plays an important role in the maintenance and improvement of a wide range of cognitive processes (Rasch and Born 2013; Mednick et al. 2011; Lowe, Safati, and Hall 2017; Whitney et al. 2017), including executive functions, (e.g., sustained attention and WM (Könen, Dirk, & Schmiedek, 2015; Vriend et al., 2013; Cellini et al. 2015; Goel et al. 2009; Lo et al. 2012)). Sleep deprivation negatively impacts WM performance as measured with digit span (Quigley, Green, Morgan, Idzikowski, & King, 2000) and the N-back task (Choo, Lee, Venkatraman, Sheu, & Chee, 2005). Additionally, studies suggest that sleep between WM training sessions, compared to wake, may be critical for enhancing WM performance (Zinke, Noack, and Born 2018; Kuriyama et al. 2008b; Lau et al. 2015). For example, training participants on an n-back task over several sessions improved accuracy of performance, but only if the interval between training sessions contained nocturnal sleep (Zinke, Noack, and Born 2018; Kuriyama et al. 2008b) or a nap (Lau et al. 2015), but not wake. Recent observations suggest that SWS may provide an optimal brain state for modification of prefrontal functioning. SWS constitutes between 10 and 25% of total sleep time (Ohayon et al. 2004) and is thought to play an important role in cerebral restoration and recovery (Horne 1992), and SWA is a global index of sleep homeostasis (Borbély and Achermann 2000). Several studies have shown a specific association between EEG activity during SWS in the enhancement of WM suggesting that SWA may reflect localized, experience-dependent, cortical plasticity (Rodriguez et al. 2016; Miyamoto, Hirai, and Murayama 2017; Huber et al. 2004). Specifically, Pugin and colleagues demonstrated frontoparietal increases in SWA following WM training, and the magnitude of SWA correlated with WM performance after three weeks of training (Pugin et al., 2015). Furthermore, SWA predicted WM gains across a period of sleep in both young (Ferrarelli et al. 2019) and older adults (Sattari et al. 2019). Taken together, these studies suggest that experience-induced changes in SWA support efficient WM improvement.

In recent years, a number of studies have explored the association between central and autonomic activity during sleep, offering promising new perspectives to understand brain-body interplay in humans (Brandenberger et al. 2001a; Thomas et al. 2014; Ako et al. 2003). For example, a recent study investigated the relationship between cardiac activity and K-complexes (KC; 0.5-1 Hz)-a positive-negative-positive waveform during Stage 2 sleep similar to slow oscillations-and demonstrated that KCs were associated with a biphasic cardiac response, with a marked heart rate acceleration followed by a gradual deceleration (de Zambotti et al. 2016). Interestingly, this biphasic fluctuation in heart rate has also been shown to coincide with bursts of K-complexes and delta waves (Sforza, Jouny, and Ibanez 2000), which, together, indicate a synchronization of central and autonomic events. Furthermore, our group recently identified cardiovascular events during NREM sleep, termed heart rate bursts, that last 2-3 seconds and occur mostly in Stage 2 and SWS (Naji et al. 2019). Heart rate bursts are temporally coincident with EEG activity, including slow oscillations and spindle/sigma activity (12-15Hz) and are followed by vagal rebound reflected in a surge in the high frequency component (HF; 0.15–0.4 Hz) of the ECG. These ***A**utonomic/**C**entral **E**vents (**ACE**s)* significantly predicted the magnitude of long-term, episodic memory improvement to a greater extent than non-ACE sleep activity and overall sleep activity. The current study builds on these findings by investigating the role of ACE activity in WM improvement.

We tested whether ACE activity during SWS sleep supports sleep-related improvement in the Operation Span Task (OSpan), a dual-task consisting of a processing subtask and a short-term memory subtask that has been commonly used to test central constructs of WM. In summary, the current study aimed: (1) to replicate ACE activity in sleep during a daytime nap; and (2) to determine the impact of ACE activity during SWS on WM gains. Given prior associations between SWA and WM improvement, we hypothesized that ACE SWA would be significantly associated with WM gains across the nap. We considered ACE sigma activity as a positive control that would not be correlated with WM improvement.

## 2. Methods

### 2.1 Participants

104 young (Age:17-23 [Mean=20.7, SD= 2.95], 64 males) healthy adults with no personal history of neurological, psychological, or other chronic illness provided informed consent, which was approved by the University of California, Riverside Human Research Review Board. Participants were randomized to either have a 2-hour nap opportunity monitored with polysomnography (PSG) (n=53), stay awake (n=51), where subjects engaged typical daily activities with actigraph monitoring. Participants included in the study had a regular sleep-wake schedule (reporting a habitual time in bed of about 7–9 h), and no presence or history of sleep, psychiatric, cardiovascular, or neurological disorder determined during an in-person, online, or telephone interview. Participants received monetary compensation for participating in the study. For PSG measures, 4 participants’ nap recordings were not collected due to recording computer failures. Among the 50 participants whose PSG were recorded, 2 of them did not have stage 2 sleep and 13 of them did not have SWS.

### 2.2. Working memory task

The current study used the Operation Span Task (OSpan; Turner & Engle, 1989; see Figure 1b) as a measure of WM performance, which requires participants to solve a series of math operations while trying to memorize a set of unrelated letters. The task was programmed in Matlab (The MathWorks Inc., 2015) using Psychtoolbox (Kleiner et al., 2007). The task included 3 practice and 20 test trials. Participants were tested in letter strings of seven. For each letter string, participants were shown a series of math problems that they had to confirm were correct within 3 seconds, using pre-determined responses on the keyboard. After each equation, a letter would appear on the screen and the subject was instructed to remember each letter. At the end of each string, the participant was instructed to recall the letters in the order presented by typing responses on a computer keyboard. Immediately after each trial, the next letter string would be presented. If the participants forgot one of the letters in a trial, they were instructed to provide their best guess. In addition, to decrease trade-off between solving the operations and remembering the letters, a 70% accuracy criterion on the math operations was required for all the participants. We excluded 2 participants based on this criterion. We calculated performance as: number of correct letters recalled in the correct order/ total number of letters in the string. For assessing change in performance from session 1 and session 2, we calculated the difference in performance between the two sessions (session 2 – session 1).

**Figure 1.**
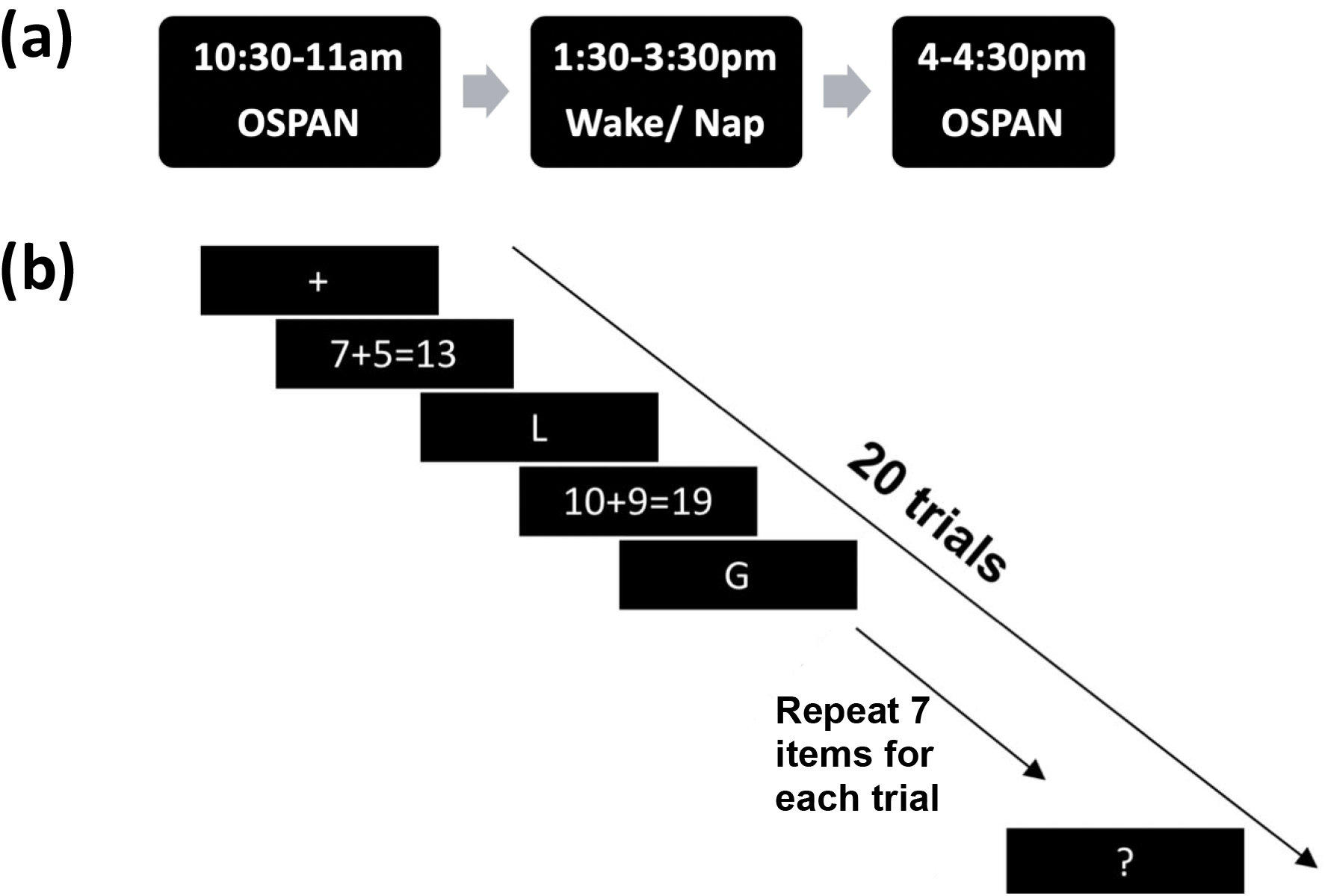
WM task (OSpan). (a) Subjects completed the 30-min OSpan WM task at 10:30 AM. Next, they were randomly assigned to either a nap condition (Nap) or a wake condition (Wake) that took place between 1:30-3:30PM. Between 4 to 4:30 PM, subjects repeated the WM task. (b) The OSpan task included 3 practice and 20 test trials. Participants were tested in letter strings seven. For each letter string, participants were shown a series of math problems that they had to confirm were correct within 3 seconds by entering responses on the keyboard. After each equation, a letter would appear on the screen and the subject was instructed to remember each letter. At the end of each string, the participant was instructed to recall the letters by typing responses on the keyboard. Immediately after each trial, the next letter string would be presented.

### 2.3. Data acquisition and pre-processing

#### 2.3.1 Study Procedure

Participants were asked to refrain from consuming caffeine, alcohol, and all stimulants for 24 h prior to and including the study day. Participants filled out sleep diaries for one week prior to the experiment and wore wrist-based activity monitors the night before the study (Actiwatch Spectrum, Philips Respironics, Bend, OR, USA) to ensure participants were well-rested, defined as at least 7 hours on average during the week and the eve of the experimental day. On the experimental day, participants arrived at the Sleep and Cognition lab at 10:00AM and had EEG electrodes attached, followed by an Operation Span (OSpan) working memory task. Nap/Wake interventions occurred between 1:30-3:30 PM. At 1:30PM, subjects in the Nap group were given 2-hours time-in-bed to obtain up to 90-min total sleep time during which their sleep were polysomnographically-recorded. Sleep was monitored online by a trained sleep technician. Subjects in the Wake group were asked not to nap, exercise, or consume caffeine or alcohol, and were monitored with actigraphy during the active wake period. All subjects were retested on the memory task between 4 and 4:30PM. Subjects completed the Karolinska Sleepiness Scale (KSS, Åkerstedt et al., 1990) questionnaire two times throughout the experimental day; at the start of each WM task (Session 1 and Session 2) to report their sleepiness. KSS is a 9-point Likert scale often used when conducting studies involving self-reported, subjective assessment of an individual’s level of drowsiness at the time, in which a higher score yields a sleepier state.

#### 2.3.2 Sleep recording and scoring

Polysomnography (PSG) data, including electroencephalogram (EEG), electrocardiogram (ECG), chin electromyogram (EMG), and electrooculogram (EOG), were collected using a 32-channel cap (EASEYCAP GmbH) with Ag/AgCI electrodes placed according to the international 10–20 System (Jasper, 1958). Electrodes included 24 scalp, two electrocardiogram (ECG), two electromyogram (EMG), two electrooculogram (EOG), 1 ground, and 1 on-line common reference channel. EEG signals were recorded at a 1000 Hz sampling rate and referenced on-line to the common reference channel. Scalp EEG and electrooculographic (EOG) electrodes were referenced to unlinked contralateral mastoids (F3/A2, F4/A1, C3/A2, C4/A1, P3/A2, P4/A1, O1/A2, O2/A1, LOC/A2, ROC/A1), and two EMG electrodes were attached under the chin and referenced to each other. After recording, all data were then digitized at 256 Hz. High-pass filters were set at 0.3 Hz, and low-pass filters were set at 35 Hz for EEG and EOG electrodes. A notch filter was set at 60 Hz. The EEG data were scored using Hume, a custom MATLAB toolbox. The records were scored in 30-second epochs using eight scalp electrodes: Frontal (F3, F4), Central (C3, C4), Parietal (P3, P4), and Occipital (O1, O2). Next, all epochs of the filtered data with artifacts and arousals were identified by visual inspection and rejected. Five sleep stages (i.e., wake, Stage 1, Stage 2, SWS, and REM) were reclassified in continuative and undisturbed 3-min bins and the bins were used for further analysis.

#### 2.3.3 Power spectral analysis

The EEG power spectrum was computed using the Welch method (4 sec Hanning windows with 50 % overlap). The frequency for sigma power was 12-15Hz and for SWA was .5–1 Hz. For RR time-series, the power spectral estimation was performed by the autoregressive model and the model order was set at 16.

### 2.4 Autonomic/Central Event Detection

#### 2.4.1 Heart-beat detection

Electrocardiogram (ECG) data were acquired at a 1000-Hz sampling rate using a modified Lead II Einthoven configuration. The ECG signals were filtered with a passband of 0.5-100 Hz by Butterworth filter. R waves were identified in the ECG using the Pan-Tompkins method, and confirmed with visual inspection. In order to extract continuous RR tachograms, the RR intervals were resampled (at 4 Hz for power spectrum estimation) and interpolated by piecewise cubic spline. Zero-phase Butterworth filters were applied to the interpolated RR time-series to extract RR_HF_.

#### 2.4.2 HR burst detection

Within 3-min bins of the RR time-series during wake and sleep stages, the HR burst events were detected as RR intervals shorter than two standard deviations below the mean.

### 2.5 Event-based Analysis for Autonomic/Central Event

#### 2.5.1 Time-locked analysis

In order to calculate changes in SWA and sigma power around the HR burst, the Hilbert transform was applied on filtered EEG signals in bands of interest (0.5–1 Hz for SWA and 12–15 Hz for sigma activity). To assess the HF amplitude fluctuation around the HR burst (RR_HF_), the Hilbert transform was applied on RR_HF_ (0.15–0.4 Hz). See Naji et al 2018 for detailed methods.

#### 2.5.2 ACE Change scores

We investigated ACE coupling during wake and sleep stages by tracking fluctuations in the EEG in a 20-sec window from 10 second before to 10 second after the HR burst peak. As we were specifically interested in sleep EEG activity previously demonstrated to correlate with WM improvement, we focused on SWA as our primary frequency band of interest and sigma activity as a positive control band. In addition, we examined RR_HF_ in the ECG channel, about which we did not have a specific hypothesis.

EEG/ECG data were binned into 5-sec intervals within the 20-sec windows around the HR burst, named −10, −5, +5, +10 window. The average RR_HF_, SWA, and sigma activity were calculated in each of the four 5-sec windows. For non-ACE brain activity, we calculate average RR_HF_, SWA, and sigma activity in periods with no HR burst (including 20 s windows around them). We computed ACE change scores for each 5-sec interval as follows: (ACE activity in each 5 s interval – non-ACE activity)/ (ACE activity in each 5 s interval + non-ACE activity). We computed similar change scores for RR_HF_. Given prior findings, we specifically focused two ACE change scores: (1) PreBase: the change score of the −5 window, and (2) PrePost: ACE activity in the −5 window subtracted from +5 window. Activity in frontal areas were averaged across F3 and F4 channels and activity in central areas were averaged across C3 and C4. Besides ACE and non-ACE activity, overall sleep values (SWA and sigma power) were calculated as average EEG power in the entire sleep stage (regardless of ACE or non-ACE).

### 2.6 Statistics Analyses

Statistical analyses were conducted using R version 3.4.3, and alpha was set at <.05.

#### 2.6.1 Event-locked analysis

For Stage 2 and SWS, repeated measures ANOVAs used to compare ACE activity across the four 5-sec bins (10-, 5-, 5+, 10+) around the HR bursts, followed by post hoc comparisons with Bonferroni corrections.

#### 2.6.2 Repeated measures ANOVAs

Repeated measures ANOVAs with two within-subject factors (four windows and two areas: frontal vs. central areas) were performed to test differences in ACE change scores across the four 5-sec bins. Greenhouse-Geisser corrections were used to adjust p-values for lack of sphericity. Post hoc comparisons were corrected using the Bonferroni method. Generalized Eta-Squared was used as a measure of effect size.

#### 2.6.3 Correlations

To investigate the contribution of ACE vs non-ACE events to WM improvement across the nap, we computed Pearson’s correlations with WM improvement.

#### 2.6.4 Regression Models

To assess the relative importance of ACE and non-ACE activity for WM improvement, we utilized a hierarchical, linear regression approach. Two linear regression models were built to predict WM gain. In Model 1, Session1 WM performance and overall frontal SWA in SWS were the independent variables. In Model 2, we added the ACE change scores (PreBase & PrePost) for frontal SWA in SWS. Baseline WM performance was included in the models as a covariate to account for individual differences in WM capacity (Matysiak, Kroemeke, and Brzezicka 2019). The regression results for the averaged frontal and central activity are tabulated in Table 1 and Table 2, respectively. By comparing Model 1 and 2, we measure the explanatory gain of ACE values over and above general sleep EEG and WM capacity for WM improvement.

**Table 1:** Change Scores for ACE variables during SWS.

| Window |  |  | -10 | -5 | +5 | +10 |
| --- | --- | --- | --- | --- | --- | --- |
| SWA | Frontal | Mean | -0.036 | 0.183 | 0.047 | 0 |
|  |  | S.E. | 0.024 | 0.034 | 0.030 | 0.030 |
|  | Central | Mean | -0.047 | 0.197 | 0.047 | 0.022 |
|  |  | S.E. | 0.167 | 0.189 | 0.175 | 0.171 |
| Sigma | Frontal | Mean | -0.006 | 0.083 | 0.014 | -0.003 |
|  |  | S.E. | 0.022 | 0.014 | 0.022 | 0.018 |
|  | Central | Mean | 0.003 | 0.072 | 0.015 | -0.005 |
|  |  | S.E. | 0.020 | 0.015 | 0.021 | 0.018 |
| $RR_{HF}$ | | Mean | 0.022 | 0.076 | 0.138 | 0.045 |
|  |  | S.E. | 0.016 | 0.022 | 0.022 | 0.022 |

**Table 2:** Change Scores for ACE variables during Stage 2 Sleep.

| Window |  |  | -10 | -5 | +5 | +10 |
| --- | --- | --- | --- | --- | --- | --- |
| SWA | Frontal | Mean | -0.036 | 0.147 | -0.026 | -0.047 |
|  |  | S.E. | 0.011 | 0.019 | 0.011 | 0.009 |
|  | Central | Mean | -0.034 | 0.132 | -0.023 | -0.040 |
|  |  | S.E. | 0.010 | 0.016 | 0.010 | 0.009 |
| Sigma | Frontal | Mean | -0.008 | 0.137 | -0.008 | -0.098 |
|  |  | S.E. | 0.017 | 0.020 | 0.016 | 0.019 |
|  | Central | Mean | -0.011 | 0.117 | -0.020 | -0.104 |
|  |  | S.E. | 0.016 | 0.017 | 0.015 | 0.019 |
| RR<sub>HF</sub> |  | Mean | 0.006 | 0.062 | 0.179 | 0.029 |
|  |  | S.E. | 0.009 | 0.014 | 0.014 | 0.010 |

## Results

### Working Memory Performance: Comparing Nap vs Wake

Our analysis revealed no significant difference in WM between the two nap conditions in Session 1 [F_(1,101)_ = 0.804, p = 0.372]. We compared differences in WM improvement after either a nap or wake period using a one-way ANOVA with Nap condition (Nap vs. Wake) as the independent variables, WM improvement as the dependent variable. The analysis revealed a main effect of nap condition [F_(1,101)_ = 5.734, p = 0.0185], in which participants showed a greater differential benefit from the nap compared to the wake condition.

### EEG power is modulated by RR phase

For SWA change scores, a significant effect of windows was found during Stage 2 (F_(3, 144)_ = 61.434, p<.001, η_p_^2^ = .418), as well as during SWS (F_(3, 111)_ = 17.307, p<.001, η_p_^2^ = .183). Post hoc comparisons revealed the highest SWA occurred during the −5 window (prior to the HR burst) during Stage 2 (all ps < 0.001), as well as during SWS (all ps < 0.001). The interactions between windows and channels was significant during Stage 2 (F_(3,144)_ = 5.645, p= .001, η_p_^2^ = .003), but not SWS (p= .350), although none of the post hoc comparisons was significant (all ps > 0.225).

For Sigma change scores, repeated measures ANOVAs also indicated significant effect of window in sigma activity change scores during Stage 2 (F_(3, 144)_ = 46.53, p<.001, η_p_^2^ = .321) and SWS (F_(3, 111)_ = 10.513, p < .001, η_p_^2^ = .078). Post hoc comparisons revealed the highest sigma activity occurred during the −5 window during Stage 2 (all ps < 0.001), as well as during SWS (all ps < 0.001). The interactions between windows and channels was significant for Stage 2 (F_(3, 144)_ = 6.097, p= .001, η_p_^2^ = .001) and SWS (F_(3, 111)_ = 3.711, p = 0.03, η_p_^2^ = .001), although none of the post hoc comparisons was significant (all ps > 0.180).

For RR_HF_, repeated measures ANOVAs with a within-subject factor (windows) revealed significant differences in RR_HF_ activity change scores during Stage 2 (F_(3, 144)_ = 57.749, p<.001, η_p_^2^ = .391), as well as during SWS (F_(3, 111)_ = 21.015, p < .001, η_p_^2^ = .107). Post hoc comparisons revealed that RR_HF_ activity significantly increased during the −5 window and reached the maximum level during the +5 window during Stage 2 (all ps < 0.001), as well as during SWS (all ps < 0.001).

Taken together, we replicated the profile of ACE activity in Naji et al., with sigma and SWA power in Stage 2 and SWS increased from overall average to a maximum level prior to the peak of the HR bursts (−5 window), and returned to average post-HR-peak, and RR_HF_ increased from average and reached the maximum level after the peak of the HR bursts (+5 window).

### Correlations with WM Improvement

As predicted, WM improvement was positively correlated with ACE SWA change scores in PreBase (Figure 4a Frontal: r= .447, p= .012; Figure 4c Central: r= .524, p= .003, as well as PrePost (Figure 4b Frontal: r= .560, p= .001; Figure 4d Central: r= .455, p= .010) during SWS. Interestingly, the correlations between WM improvement and overall SWA power in SWS were not significant (see Figure 5; all ps > .668). Additionally, as predicted, we found no significant correlations between WM improvement and ACE sigma change scores (PreBase: all ps > .153; PrePost: all ps > .051) or overall sigma power (all ps > .237) during SWS. For Stage 2, no significant correlations between WM improvement and ACE SWA power were found for PreBase (all ps > .439) or PrePost (all ps > .619). Similarly, we found no significant correlations between WM improvement and ACE sigma change scores (PreBase: all ps > .672; PrePost: all ps > .257) or overall sigma power (all ps > .352) during Stage 2 sleep. Lastly, we found no significant correlations between WM improvement and RR_HF_ activity during Stage 2 sleep (p = .054) or SWS (p = .392).

**Figure 2.**
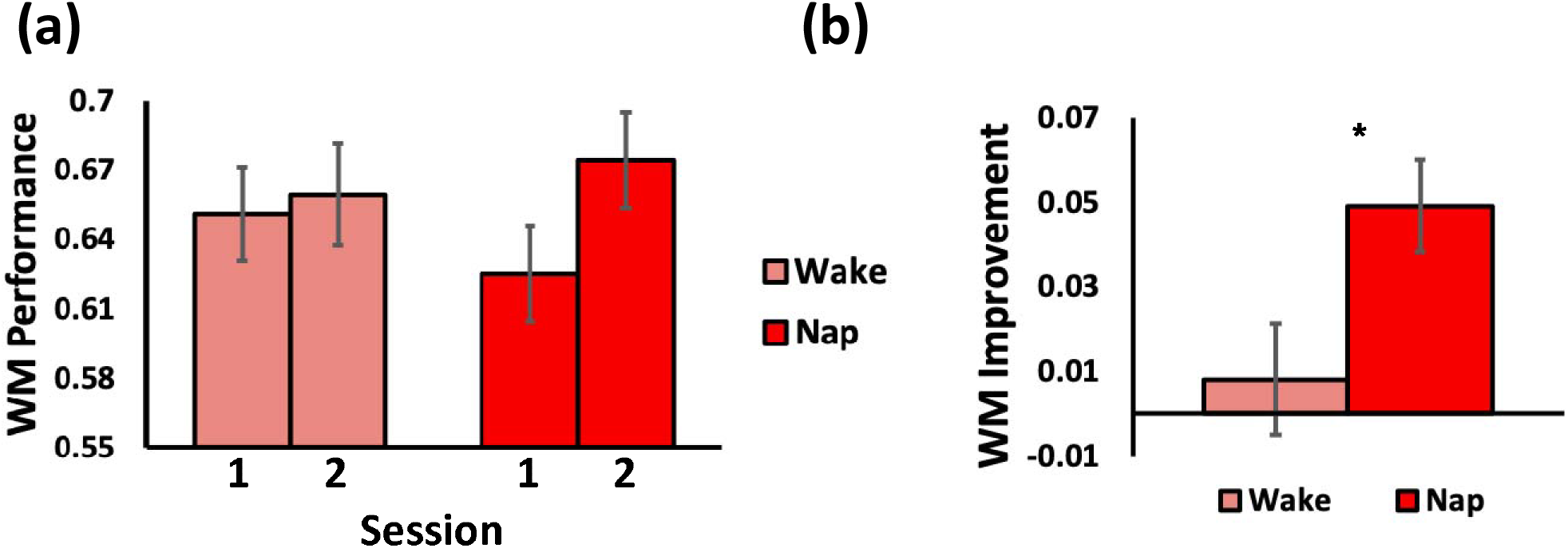
(a–b) Working Memory Performance by Nap Condition: (a) Session 1 and 2 performances by Nap Condition (b) WM improvement by Nap Condition: Significant difference in WM improvement after a nap or a period of wake was observed. Asterisks between error bars indicate significant differences between nap conditions (*p < 0.05). Error bars represent standard error of the mean.

**Figure 3:**
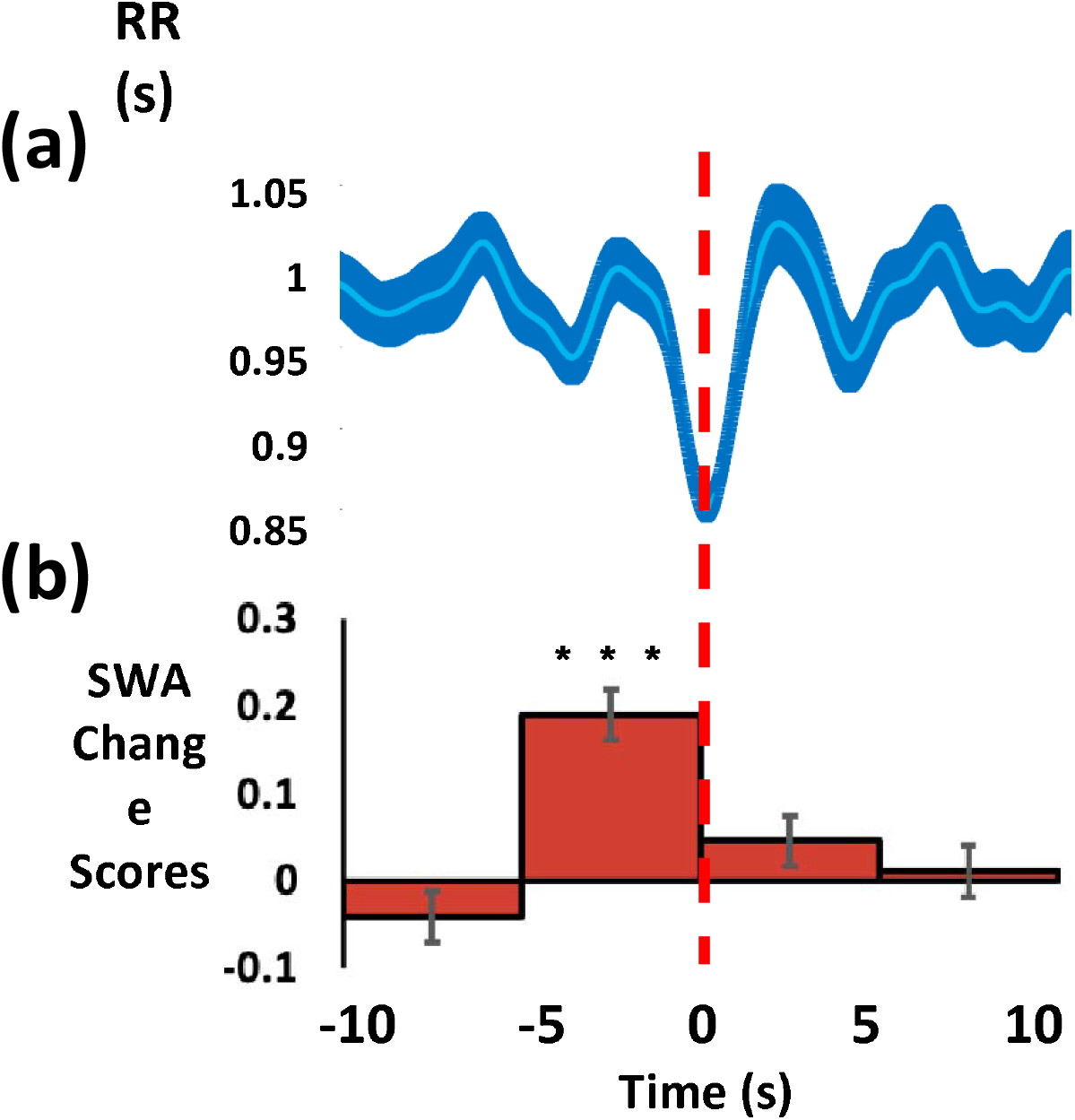
Heart-rate-bursts and SWA change scores time-locked on HR bursts during SWS. (a) Grand average of the HR bursts (b) ACE SWA change scores show a significant increase in the −5 window. Asterisks indicate significant differences from the non-ACE activity after Bonferroni correction for multiple comparisons (***p<.001).

**Figure 4:**
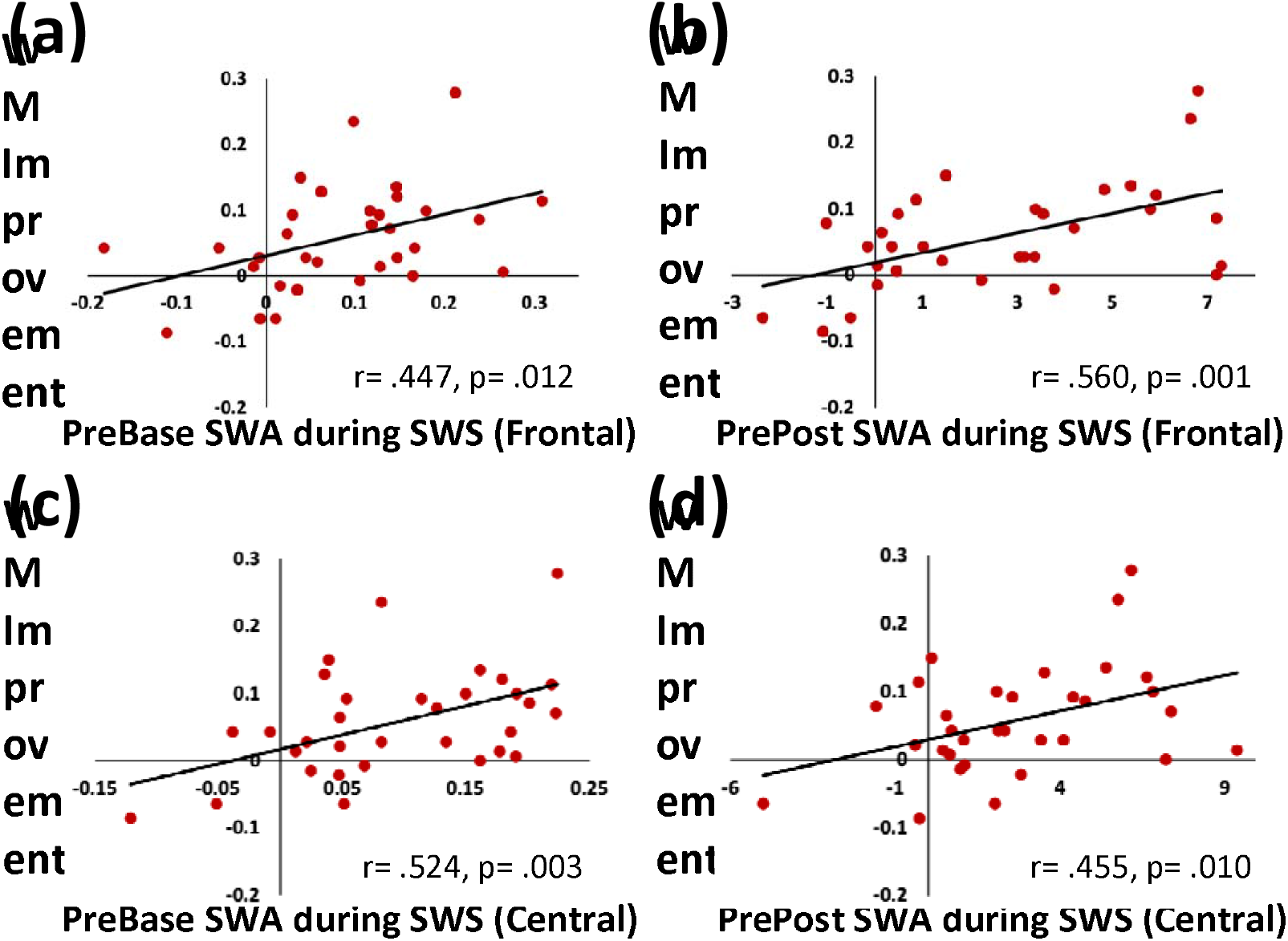
Correlations between ACE SWA change scores (X-axis) and WM improvement (Y-axis) Association between WM improvement and (a) Frontal ACE PreBase SWA during SWS (r= .447, p= .012); (b) Frontal ACE PrePost SWA during SWS (r= .560, p= .001); (c) Central ACE PreBase SWA during SWS (r= .524, p= .003); (d) Central ACE PrePost SWA during SWS (r= .455, p= .010).

**Figure 5:**
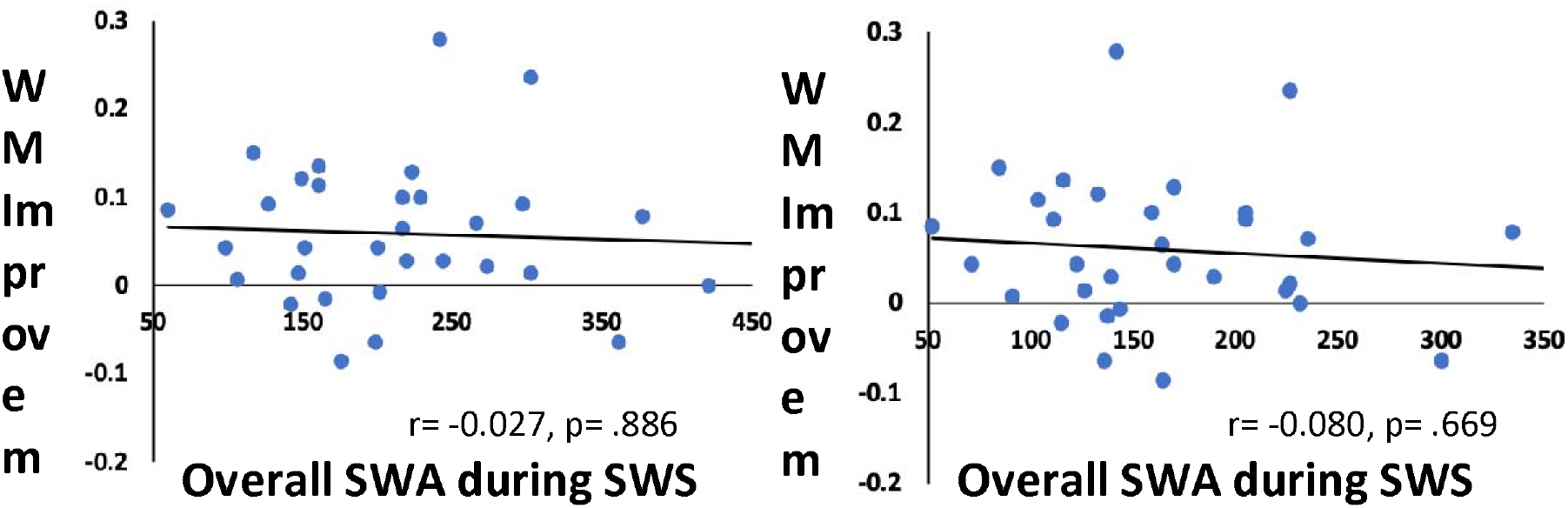
Correlations between overall SWA power (X-axis) and WM improvement (Y-axis) Association between WM improvement and (a) Frontal SWA during SWS (r= −0.027, p= .886); and (b) Central SWA during SWS (r= −0.080, p= .669);

### Importance of ACEs for WM Improvement

Next, we assessed the relative importance for WM performance of ACE and non-ACE activity using hierarchical, linear regression. The regression results using frontal and central channels are shown in Table 3 and Table 4, respectively.

**Table 3:**
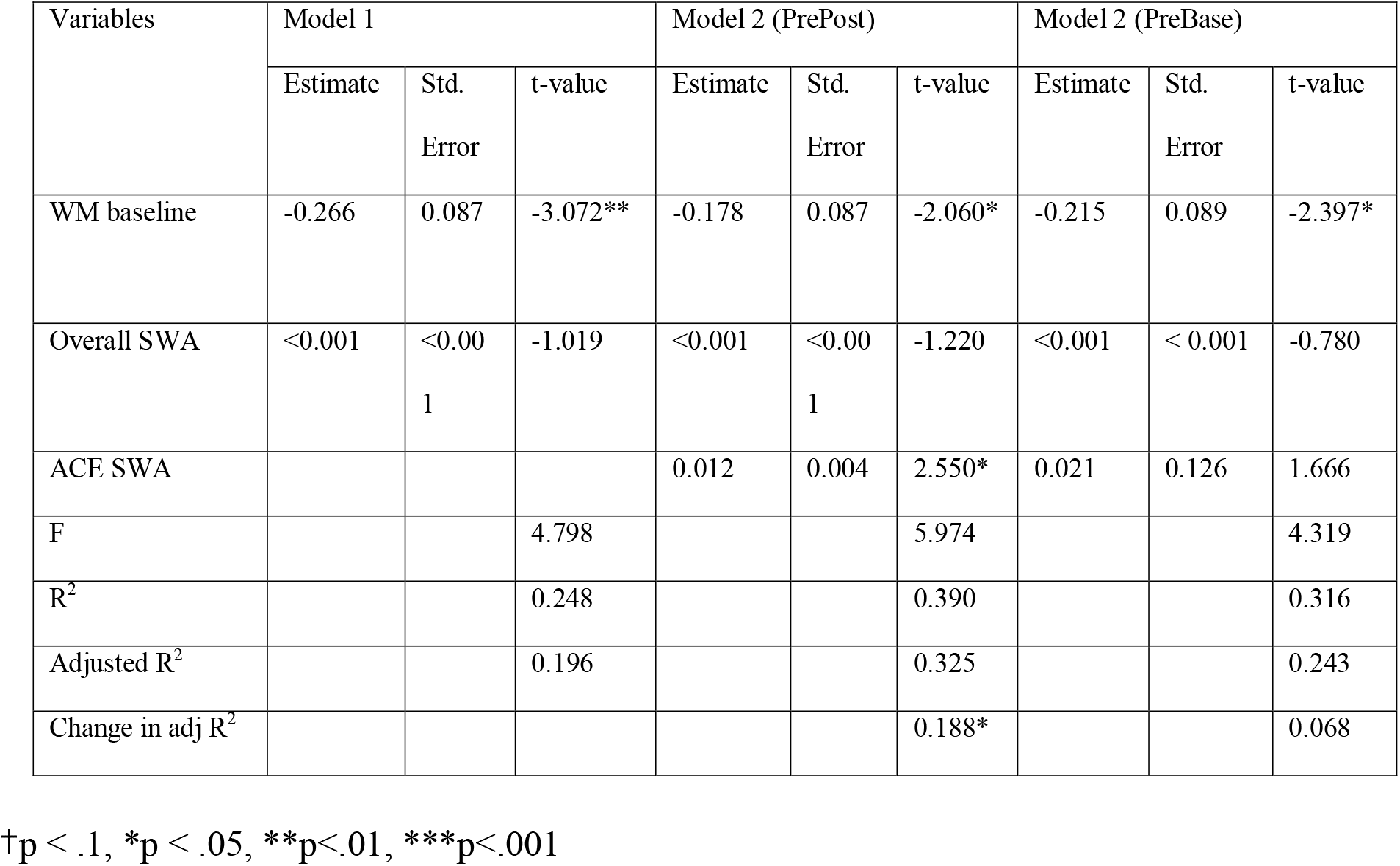
Regression models: ACE frontal channels predicting WM improvement

**Table 4:**
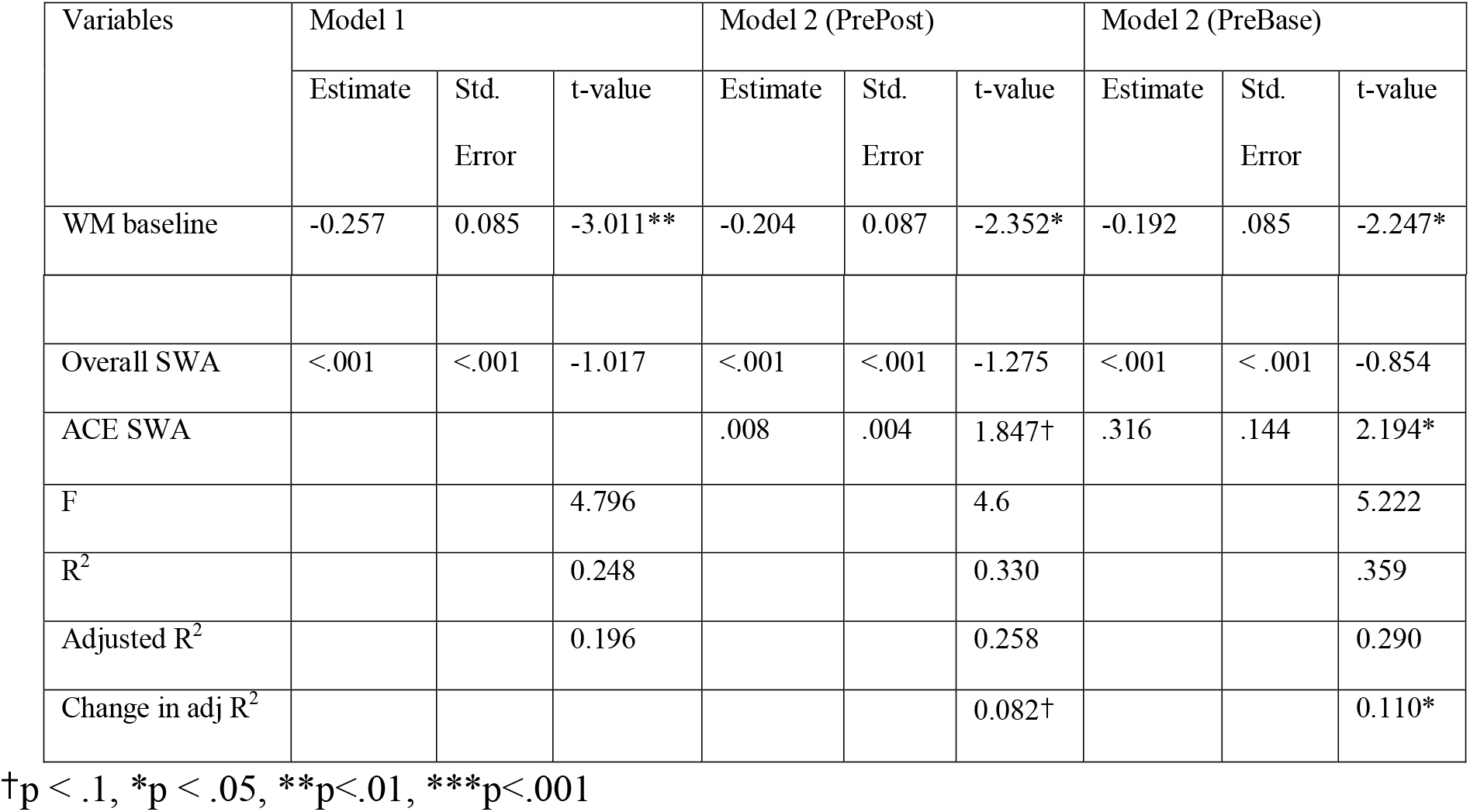
Regression models: ACE central channels predicting WM improvement

| Variables | Model 1 |  |  | Model 2 (PrePost) |  |  | Model 2 (PreBase) |  |  |
| --- | --- | --- | --- | --- | --- | --- | --- | --- | --- |
|  | Estimate | Std. Error | t-value | Estimate | Std. Error | t-value | Estimate | Std. Error | t-value |
| WM baseline | -0.257 | 0.085 | -3.011** | -0.204 | 0.087 | -2.352* | -0.192 | .085 | -2.247* |
| Overall SWA | <.001 | <.001 | -1.017 | <.001 | <.001 | -1.275 | <.001 | <.001 | -0.854 |
| ACE SWA |  |  |  | .008 | .004 | 1.847† | .316 | .144 | 2.194* |
| F |  |  | 4.796 |  |  | 4.6 |  |  | 5.222 |
| R<sup>2</sup> |  |  | 0.248 |  |  | 0.330 |  |  | .359 |
| Adjusted R<sup>2</sup> |  |  | 0.196 |  |  | 0.258 |  |  | 0.290 |
| Change in adj R<sup>2</sup> |  |  |  |  |  | 0.082† |  |  | 0.110* |

†p < .1, \*p < .05, \*\*p<.01, \*\*\*p<.001

### Regression Models using the Averaged Frontal Channel

In Model 1, WM gain was the dependent variable and Session 1 WM performance and overall frontal SWA in SWS were the independent variables. Model 1 reached significance (F_(2,29)_= 4.798, p= .016; adj R^2^ = .196), but adding the ACE measures (PrePost) in Model 2 enhanced the prediction of the model (F_(3,28)_= 5.974, p= .021; adj R^2^ = .325), with the ACE SWA (PrePost) being a significant predictor (p= .016). Model 2 accounted for significantly more of the variance in WM improvement than Model 1 (change in adj R^2^ = .142, F_(1,28)_= 6.504, p=.016). Similarly, adding the ACE SWA (PreBase) measures in Model 2 enhanced the prediction of the model (F_(3,28)_= 4.319, p= .013; adj R^2^ = .243), although non-significantly (change in adj R^2^ = .068, F_(1,28)_= 2.774, p=.106).

### Regression Models using the Averaged Central Channel

In Model 1, baseline WM performance and overall central SWA in SWS were the independent variables. Model 1 reached significance (F_(2,29)_= 4.796, p= .016; adj R = .196), but adding the ACE SWA measures (PreBase) in Model 2 elevated significance of the model (F_(3,28)_= 5.222, p= .005; adj R^2^ = .290), with the ACE SWA (PreBase) being a significant predictor (p= .036). Again, Model 2 accounted for significantly more of the variance in WM improvement than Model 1 (change in adj R^2^ = .147, F_(1,28)_= 4.812, p=.036). Similarly, adding the ACE SWA measures (PrePost) in Model 2 elevated the model (F_(3,28)_= 4.6, p= .009; adj R^2^ = .258), although non-significantly (change in adj R^2^ = .109, F_(1,28)_= 3.411, p=.075).

In summary, while the overall SWA power did not predict individual differences in sleep-dependent WM gain, ACE events predicted up to 32.5% of the variance in performance improvement on this WM task.

## Discussion

Consistent with Naji et.al (2017), we confirmed an autonomic cardiac event during NREM sleep that is temporally-coupled with a significant boost in brain oscillations, and reported that these autonomic/central events (ACEs) contribute to WM improvement. Specifically, we showed that increases in the EEG amplitude in SWA and sigma bands preceded the large-amplitude HR bursts. Furthermore, we showed that these time-locked boosts in SWA during SWS, but not sigma or RR_HF_, can predict WM improvement across a daytime nap to a greater extent than overall SWA. Taken together, the results suggest that heart-brain interactions during sleep may be a critical mechanism for sleep-related WM gain.

### Heart-brain interaction during sleep: findings and potential mechanisms

Emergent research examining brain-body communication suggests that autonomic activity may be linked with sleep brain activity, and that this interaction is likely a distinct predictor of plasticity, cognitive ability and enhancement. Studies have revealed a consistent symmetry between heart and brain activity with temporal changes in NREM delta (0.5-4Hz) power and ANS activity (Brandenberger et al. 2001b; Yang et al. 2002; Kuo and Yang 2004; Ako et al. 2003; Rothenberger et al. 2015; Jurysta et al. 2003; 2005; Thomas et al. 2014). Delta EEG power, a marker of homeostatic sleep drive dissipates across successive NREM periods. Brandenberger and colleagues (2001) demonstrated an inverse coupling between oscillations in delta wave activity (0.5-3.5Hz) and autonomic activity during nighttime sleep. Using a cross-correlation approach, Thomas et al. (2014) showed a temporal relationship between SWA and high frequency cardiopulmonary (0.1-0.4Hz) coupling, an ECG-derived biomarker of stable sleep, during both stage 2 sleep and SWS. These findings suggest that CNS-ANS dynamics support the interdependency between cortical and cardiac function during sleep.

Several studies have attempted to provide insight on directionality and potential mechanisms of heart-brain communication. Using spectral coherence analysis, Jurysta and colleagues reported that alterations in cardiac parasympathetic activity virtually paralleled changes in EEG delta power, with parasympathetic activity consistently preceding EEG delta changes by about 12 min. This suggested that increases in vagal modulation may precede increases in EEG delta power (Jurysta et al. 2003; 2005). On the other hand, splitting a night into four NREM sleep cycles, Rothenberger and colleagues showed that the time-varying correlations between delta EEG power and heart rate variability (HRV) were largest in the first NREM period, followed by the second and third (Rothenberger et al. 2015). Using this approach, the EEG-HRV coupling appeared to reflect the dynamics of delta, which suggests that the relationship may be more strongly driven by CNS (and potentially by homeostatic sleep factors) than ANS activities. Although more research is needed to understand the complicated relation between the ANS and CNS, one possible reason for the discrepancy regarding ANS-CNS coupling in previous work may be due to the method of averaging across large periods of the night that underestimate tight temporal interactions between the heart and brain. Traditional measures examine activity over 5minute windows, which may average out fluctuations that occur in shorter time scales (e.g., heart rate bursts <5 sec). Building upon these findings, the current study adopted a high temporal precision time-domain analysis approach to the cardiac signal, which allowed for the identification of HR burst that lasted 4-5 seconds and predominated (~1 per minute) in NREM sleep. By focusing on beat-to-beat changes in sleep ECG/EEG, the current study attempted to demonstrate functional significance of ACE activity for cognitive processes.

While mechanisms driving EEG fluctuations time-locked on HR bursts remains unclear, some evidence points to arousal responses during sleep. In line with our findings, de Zambotti et al. (2016) showed that tone-triggered K-complexes are temporally coupled with a rapid increase and then decrease in heart rate activity, and coincide with bursts of K-complexes and slow waves (Sforza, Jouny, and Ibanez 2000). In this context, both synchronous EEG (K-complexes, or bursts of SOs) and cardiovascular activations (heart-rate acceleration) were viewed as responses to arousal from sleep, as when acoustic tones were not accompanied by a K-complex, the heart rate fluctuation was reduced or absent, indicating that arousal responses might be driving ANS/CNS activities. It’s been hypothesized that in the case of the K-complex, the recruited synchronized EEG response acts as a mechanism to decrease cortical arousal, suggesting that the heart-rate acceleration (tachycardia, or HR bursts) can be viewed as peripheral response of arousal from sleep. Taken together, the subsequent heart-rate deceleration that de Zambotti et al. (2016) showed and the surge of HF that Naji et al. (2019) and the current study found may reflect a feedback effect of arousal showing an inertial effect once the arousal stimulus is removed. Alternatively, arousal and post-arousal periods may modulate the autonomic system reflecting the activation-deactivation of neuronal oscillations intrinsically regulated by the cyclic arousability of the sleepy brain (Schnall et al., 1999; Sforza et al., 1999; Ferri et al., 2000).

In addition to arousal responses, ANS-CNS coupling events may also represent the brainstem’s dynamic maintenance of homeostasis during the transition from wake to sleep. As homeostatic pressure drives the transition from lighter sleep to deeper stages, the CNS and ANS experience large and rapid slowing in physiological rhythms, including decreased heart rate, broad synchronization of EEG slow waves (Fernandez Guerrero and Achermann 2018), as well as alignment of cortical and autonomic signals (Ulke et al. 2017). Brainstem medullary nuclei are responsible for a wide range of bodily functions including deepening of slow wave sleep (Anaclet and Fuller 2017) and deceleration of heart rate (Monge Argilés et al. 2000). It is, therefore, possible that medullary nuclei regulating wake to sleep transitions modulate the pace of the slow down with brief accelerations in cardiac activity. Due to the relative overlap between nuclei, these fluctuations may promote time-locked ACEs between sleep promoting oscillations such as SOs and spindles. Thus, ACE coupling may represent the adaptive and flexible modulation of central and peripheral activities by the brainstem. Given this hypothesis, we predict that brain areas involved in ACE activity are overlapping and simultaneously activate cardiac changes and initiate SWA and spindles. In this framework, these cognition-related EEG events may increase as a side-effect of homeostasis, a speculation that has interesting implications for aging research and evolutionary theory. However, more research is needed to make definite conclusions. Regardless of mechanism, these interactions have an intriguing interplay with sleep-related cognitive plasticity.

### Heart-brain interactions during sleep for cognition

Though the functional significance of CNS-ANS couplings during sleep is only beginning to be explored, we hypothesize that ACE activity may play an important role in hippocampal-prefrontal cognitive enhancement. Results from our group have shown ACE (SWA and spindles) contributions to long-term, episodic memory (Naji et al. 2019), as well as WM gains (SWA, current study). In addition, recent studies implicate frontal SWA with improvement in WM, a cognitive ability strongly supported by the prefrontal cortex (Wager and Smith 2003). For example, Ferrarelli et al. (2019) demonstrated that fronto-parietal SWA during nocturnal sleep can predict the WM improvement across the sleep. Similarly, Sattari et al. (2019) showed that frontal SWA, but not sigma, during a nap predicted WM improvement in older adults. Furthermore, ventromedial prefrontal cortex has been shown to regulate both vagal activity and slow oscillations (Dang-Vu et al., 2010; Thayer & Lane, 2000), and higher waking vagal activity is associated with better executive function (including WM). Anatomically, bidirectional projections from the prefrontal cortex to the hypothalamus and brainstem create a feedback loop for communication between peripheral sites and central cognitive areas (Thayer and Lane 2009; Shaffer, McCraty, and Zerr 2014). Furthermore, the prefrontal cortex is implicated in top-down control of the vagus nerve, and prefrontal cortical thickness is positively associated with vagally-mediated autonomic activity during wake in both young (Winkelmann et al. 2016) and older adults (Lin et al. 2017). Together, these studies point to a significant role of prefrontal processing in the improvement of working and long-term memory during sleep that is mediated by autonomic activity.

The current study is the first, to our knowledge, to identify a functional role of ANS-coupled EEG fluctuations on WM improvement. Moreover, we find a high degree of specificity with ACE-SWA but not ACE-sigma or ACE-RR_HF_ benefitting performance. While some previous studies showed an association between SWA power during sleep and WM improvement (Sattari et al. 2019; Ferrarelli et al. 2019; Pugin et al. 2015), we and others (Lau et al. 2015; MacDonald et al. 2018) failed to find a correlation between WM improvement and overall SWA. One possible reason for the discrepancy may be due to the method of averaging across large periods of data that underestimate subtle EEG changes that are time-locked on HR bursts. An intriguing idea that deserves further investigation is whether these discrepancies might be explained by considering ACE activity as a moderator. We suggest that positive findings of an association between SWA and WM may have had greater ACE activity masquerading as higher overall SWA, whereas studies that showed no correlations may have had poor ACE coupling and low SWA. Adding ACE analysis to future studies will be critical for understanding potential sleep mechanisms for WM improvement.

Our data suggest that robust coupling between frontal SWA and HR reflects increased functioning of prefrontal cortex during NREM sleep, including benefits to WM and long-term memory. This hypothesis is consistent with the neurovisceral integration model (Thayer and Lane, 2000) that contends that medial prefrontal cortex regulates autonomic activity through its connections with the nucleus tractus solitarii (NTS), and proposes that autonomic activity reflects the functional capacity of the brain structures that support WM and physiological selfregulation (Thayer et al. 2009). These findings suggest the intriguing possibility that modulation of autonomic activity during sleep may provide a novel method for boosting executive function.

Vagal nerve stimulation, for example, has been considered a valuable therapeutic option for neurologic diseases, and studies have demonstrated the ability of vagal nerve stimulation during wake to modulate vagal afferents activation (Nonis et al. 2017), and to enhance verbal memory performance (Clark et al. 1999), cognitive flexibility (Ghacibeh et al. 2006), and recently, WM (Sun et al. 2017). Future research should investigate the potential benefit of sleep-related interventions (i.e. non-invasive brain stimulation and vagal nerve stimulation) on heart-brain communication and its potential benefit for cognitive enhancement or as a clinical intervention in age-related cognitive decline.

